# A Shared Amyloid Architecture in Cardiac Fibrils from Three Neuropathy-Associated ATTR Variants

**DOI:** 10.1101/2025.08.05.667944

**Authors:** Maria del Carmen Fernandez-Ramirez, Shumaila Afrin, Binh An Nguyen, Virender Singh, Maja Pekala, Preeti Singh, Yasmin Ahmed, Rose Pedretti, Andrew Lemoff, Barbara Kluve-Beckerman, Farzeen Chhapra, David Eisenberg, Lorena Saelices

**Affiliations:** Center for Alzheimer’s and Neurodegenerative Diseases, Department of Biophysics, Peter O’Donnell Jr Brain Institute, University of Texas Southwestern Medical Center (UTSW), Dallas, TX, USA; SciKonnect and BioPatriKa, India; Department of Biochemistry, University of Texas Southwestern Medical Center, Dallas, TX, USA; Department of Pathology and Laboratory Medicine, Indiana University School of Medicine, Indianapolis, IN, USA; Department of Biological Chemistry, University of California, Los Angeles, Howard Hughes Medical Institute, Los Angeles, CA, USA

**Keywords:** Transthyretin, ATTRv, amyloidosis, amyloid, transthyretin amyloid, cryoEM, cardiomyopathy, polyneuropathy, polymorphism

## Abstract

ATTR amyloidosis results from the systemic accumulation of wild-type (ATTRwt) or mutant (ATTRv) transthyretin amyloids, leading to multi-organ dysfunction and death. The disease exhibits variable pathology and penetrance, and its relationship with the amyloid structure remains unclear. Patients carrying the neuropathy-associated variants ATTRvI84S and ATTRv-V122Δ present polymorphic ATTR fibrils, in contrast to the consistent morphology reported for most ATTR fibrils to date. Here, we aim to elucidate a potential link between neuropathic symptomatology, distinct mutations, and amyloid structural diversity, using cryo-EM. We determined the *ex-vivo* fibril structures from the variants ATTRv-P24S, ATTRv-A25S, and ATTRv-D38A, whose patients presented variable clinical manifestations, including neuropathy. Our findings revealed that, despite differences in mutations and diverse clinical phenotypes, these variants share a common amyloid core previously identified in ATTRwt and several other cardiac ATTRv. This structural consistency is significant for the development of structure-guided diagnostic tools capable of addressing the diverse spectrum of ATTR amyloidosis.

**Highlights:**

- Determines transthyretin amyloid structures of three human ATTRv by cryo-EM
- Determines the structure of three ex-vivo ATTRv fibrils linked to polyneuropathy.
- Reveals structural similarities of ATTRv amyloid cores.
- Reveals a common fold despite the different mutations and symptomatology.
- Contributes to the understanding of transthyretin aggregation in patients with diverse phenotypes

## Introduction

ATTR amyloidosis is a clinically heterogeneous and degenerative disease resulting from the extracellular accumulation of amyloidogenic transthyretin (ATTR) in the form of amyloid fibrils. ATTR amyloid deposition causes multiorgan clinical manifestations, including cardiomyopathy, polyneuropathy, carpal tunnel syndrome, gastrointestinal symptoms, and renal dysfunction, among others ^1, 2^. Cardiomyopathy is the primary cause of death in wild-type ATTR (ATTRwt) amyloidosis patients ^3^. In contrast, over 200 pathogenic mutations in the *TTR* gene can give rise to variant ATTR (ATTRv) amyloidosis, which is linked to a broader and less predictable range of symptoms beyond cardiomyopathy ^4-6^. An additional challenge arises from the fact that individuals carrying the same genetic mutation can exhibit markedly different clinical symptoms, complicating both diagnosis and treatment ^7^. The molecular mechanisms underlying this phenotypic variability remain poorly understood ^8^.

Recent developments in cryo-electron microscopy (cryo-EM) have offered valuable insights into the structural characteristics of amyloid fibrils in amyloid diseases ^9-15^. For instance, in ATTRwt amyloidosis patients, our research, alongside others’, has revealed that cardiac ATTRwt fibrils exhibit consistent molecular composition and structural conformation ^16, 17^. These fibrils are composed of two main fragments: an N-terminal fragment spanning from Pro 11/Leu 12 to Arg 34/Lys 35, and a C-terminal fragment spanning from Gly 57/Leu 58 to Thr 123/Asn 124. Both fragments consist of multiple β-strands, with residues from Gly 57 to Ser 84 forming a polar channel within the C-terminal fragment. In ATTRv amyloidosis, structures of cardiac fibrils from patients with the genotypes ATTRv-V20I, ATTRv-V30M, ATTRv-G47E, ATTRv-T60A, and ATTRv-V122I also share a similar conformation ATTRwt fibrils ^18-20^. These patients typically present with cardiomyopathy or a combination of cardiomyopathy and polyneuropathy. However, our studies on cardiac fibrils from primarily polyneuropathic ATTRv amyloidosis cases, such as ATTRv-I84S or ATTRv-V122Δ heterozygous patients, have revealed additional fibril polymorphism ^9, 21^. The ATTRv-I84S fibrils display local structural variations in the fragment comprising residues Gly 57 to Gly 67, corresponding to the gate area of the polar channel ^9^. The ATTRv-V122Δ fibrils are comprised of one or two protofilaments, with the double protofilaments arranged in two distinct types of interfaces ^21^. Whether these structural differences are associated with phenotypic variation between cardiomyopathy and polyneuropathy in ATTRv patients requires further investigation.

Here, we set out to determine the cardiac fibril structures from three ATTRv genotypes— ATTRv-P24S, ATTRv-A25S, and ATTRv-D38A—associated with polyneuropathy with or without cardiomyopathy. By interrogating the ultrastructure of cardiac deposits from these patients, we aim to explore whether differences in fibril architecture could be associated with neuropathic clinical presentations, laying the groundwork for deeper investigation into structure–phenotype relationships in ATTRv amyloidosis.

## Results

### Fibril extraction, typing, and mass spectrometry analysis of ATTRv fibrils

We successfully extracted the amyloid fibrils from the hearts of three heterozygous patients with ATTRv-P24S, ATTRv-A25S, and ATTRv-D38A amyloidosis using a previously published water-based extraction protocol. ^9^ We used an in-house antibody directed against the C-terminal fragment of transthyretin to confirm the type of ATTR fibrils. ATTR amyloidosis is categorized into two types: Type A fibrils, which contain a mixture of full-length and fragmented transthyretin proteins, and Type B fibrils, composed predominantly of full-length transthyretin. These fibril types have distinct diagnostic and prognostic implications and are associated with different clinical phenotypes.^22^ Western blot analysis showed bands corresponding to fragmented ATTR (∼10-12 kDa), confirming the presence of type A fibrils in all three samples **(Figure 1A)**.

**Figure 1.**
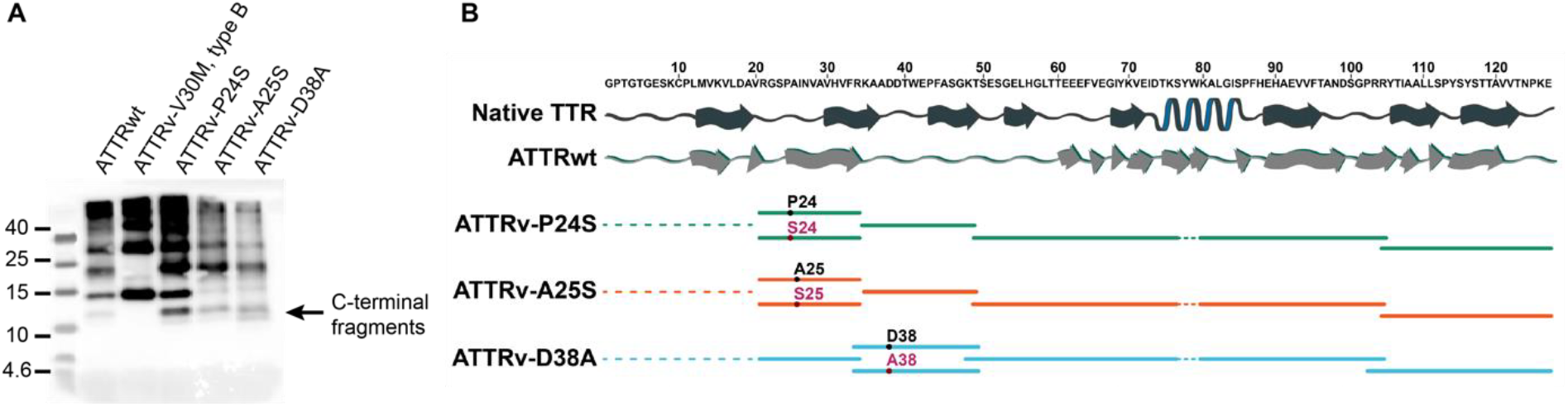
ATTRv fibrils isolated from the left ventricle of patients with different mutations. (A) Western blot of ATTRv fibrils probed with an antibody directed against the C-terminal fragment of ATTR. ATTRwt and early-onset ATTRv-V30M fibrils were used as control sample for type A and type B fibrils, respectively.^23^ (B) Mass spectrometry analysis of ATTRv-P24S, ATTRv-A25S, and ATTRv-D38A confirms the sequence identity of transthyretin consisting of both wild-type (black dots) and variant (red dots). Native sequence and structure of transthyretin are shown in black (PDB 4tlt) and the secondary structure of ATTRwt (PDB 8e7d) are shown in grey.

Mass spectrometry analysis confirmed that all three fibril samples contained both wild-type and mutant transthyretin, in agreement with the heterozygosity of the patients **(Figure 1B and Supplementary Table 2)**. Tryptic and semi-tryptic peptide analysis revealed a 1:1 wild-type to mutant ratio for ATTRv-P24S and approximately 2:1 for ATTRv-A25S and ATTRv-D38A **(Supplementary Table 3)**. The semi-tryptic results further identified the majority of enzymatic cleavage sites within the Phe44 to His56 sequence, generating C-terminal fragments characteristic of type A fibrils, which is consistent with western blot results **(Figure 1A)**.

### ATTRv-P24S, ATTRv-A25S, and ATTRv-D38A fibrils are structurally homogenous

The sample purity and concentration of extracted fibrils were evaluated by negative stain transmission electron microscopy (TEM) **(Figure 2A)**. Cryo-EM grids were prepared and screened for optimal fibril distribution prior to data-collection. The fibrils from all patients appeared to be morphologically uniform, and the fibril concentration and distribution were adequate for manual picking and helical reconstruction **(Figure 3A)**. Cryo-EM data were processed using RELION 3.1 (ATTRv-P24S) and RELION 4.0 (ATTRv-A25S and ATTRv-D38A). Two-dimensional class averages of manually picked fibrils resulted in two fibril classes, one with a clear twist, and an additional minor class of straight filaments that were not suitable for helical reconstruction **(Figure 2B)**. We used the most prevalent class of twisted fibrils for further data processing and structural determination.

**Figure 2.**
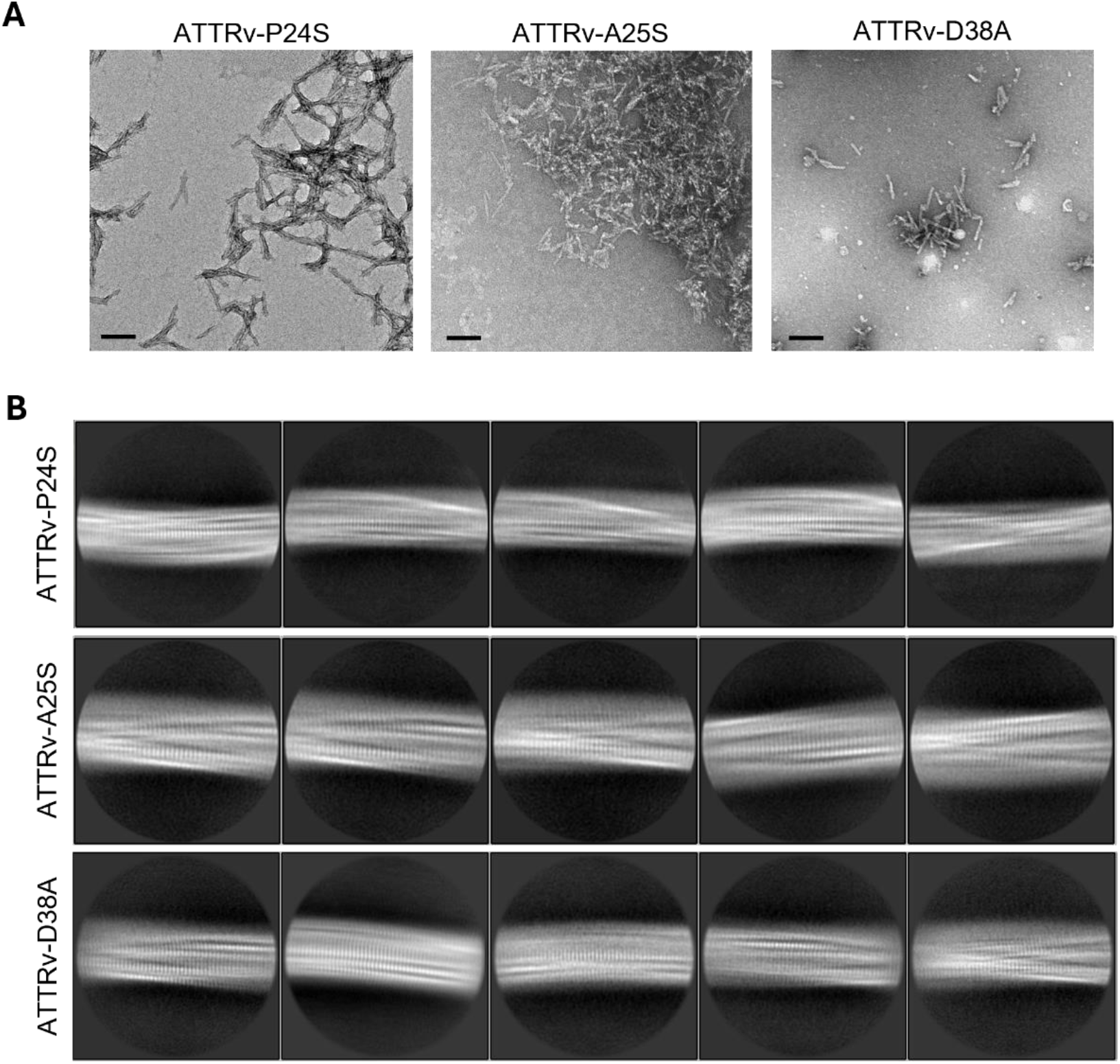
TEM images and 2D classification of the analyzed fibrils. (A) Negative stained fibrils extracted for the heart of the patients ATTRvP24S (fresh frozen tissue), ATTRvA25S and ATTRvD38A (Lyophilized tissue), visualized by transmission electron microscopy. Scale bar, 100 nm (B) Representative 2D classes obtained during the data processing.

**Figure 3.**
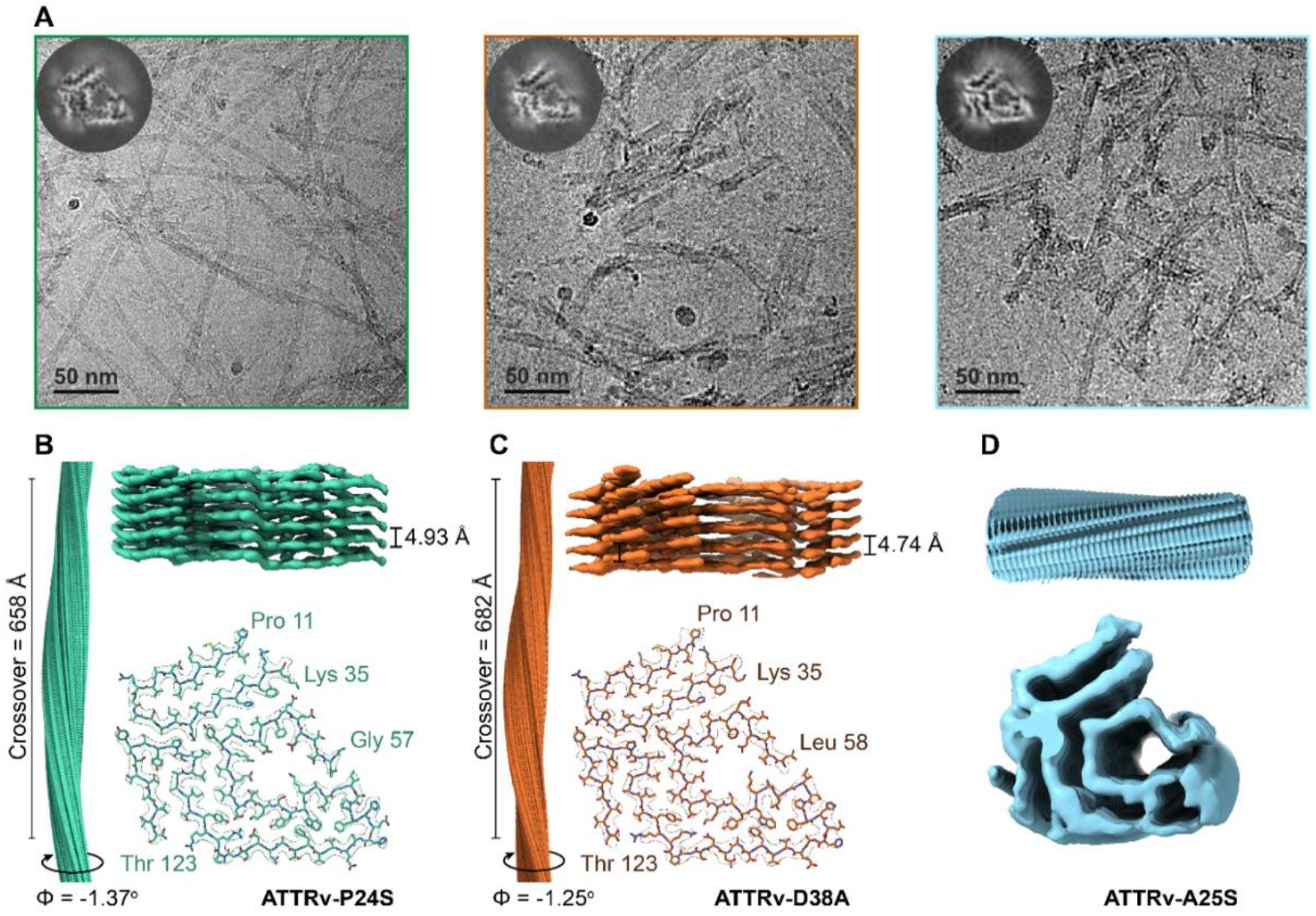
Cryo-EM images and structural analysis of ATTRv fibrils from heart. (A) Images from cryo-EM data collection of ATTRv-P24S (left), ATTRv-A25S (middle) and ATTRv-D38A (right); the major class from 3D classification of helical reconstruction is shown at the top-left corner of each image. Scale bar, 50 nm. (B-D) Cryo-EM density map and its model obtained from helical reconstruction, depicting the full crossover, twisted angle (Φ), layer separation and a model built into a single layer map. (B) ATTRv-P24S, (C) ATTRv-D38A and (D) ATTRv-A25S. The model, crossover and layer separation are not determined for ATTRv-A25S due to poor density map resolution.

The three-dimensional (3D) classification for twisted fibrils from all three samples resulted in only one 3D class **(Figure 3A)**. Aside from the minor straight filament classes that could not be reconstructed, no alternative 3D fibril morphologies were found in the present datasets. After multiple rounds of 3D refinements and local symmetry optimization, post-processing of the final density map for ATTRv-P24S and ATTRv-D38A gave the overall resolution of 3.3 Å (local resolution ranging from 3.19 – 3.81 Å) and 3.6 Å (local resolution ranging from 3.53 – 4.25 Å), respectively **(Figure 4A-D)**. ATTRv-P24S and ATTRv-D38A fibrils showed single protofilaments with layer separation and crossover distances of 4.93 Å and 658 Å, and 4.74 Å and 682 Å, respectively **(Figure 3B, C)**. The helical reconstruction of ATTRv-A25S dataset was resolved at resolution of 4.6 Å. Although we could not refine separate individual fibril layers, the cross-sectional top view of their density maps was consistent to those from ATTRv-P24S and ATTRv-D38A fibrils **(Figure 3C)**. All three structures adopt the “closed-gate” conformation, similar to previously reported structures observed in all extracts of ATTRwt and ATTRv cardiac fibrils.^16^ Similarly, the structure of ATTRv-P24S and ATTRv-D38A fibrils are composed of an N-terminal fragment spanning residues Pro 11 to Lys 35 and a C-terminal fragment spanning residues Leu 57 or Leu 58 to Thr 123 **(Figure 3B, C)**. All structures showed the presence of the polar channel formed by the C-terminal fragment enclosing residues Leu 58 to Ser 84. Residues from 1-10 and 36-56 were disordered or not present. The fuzzy white area observed around the fibril core in the cross-sectional top view of their density maps could correspond to unstructured residues from these regions **(Figure 3A)**.

**Figure 4.**
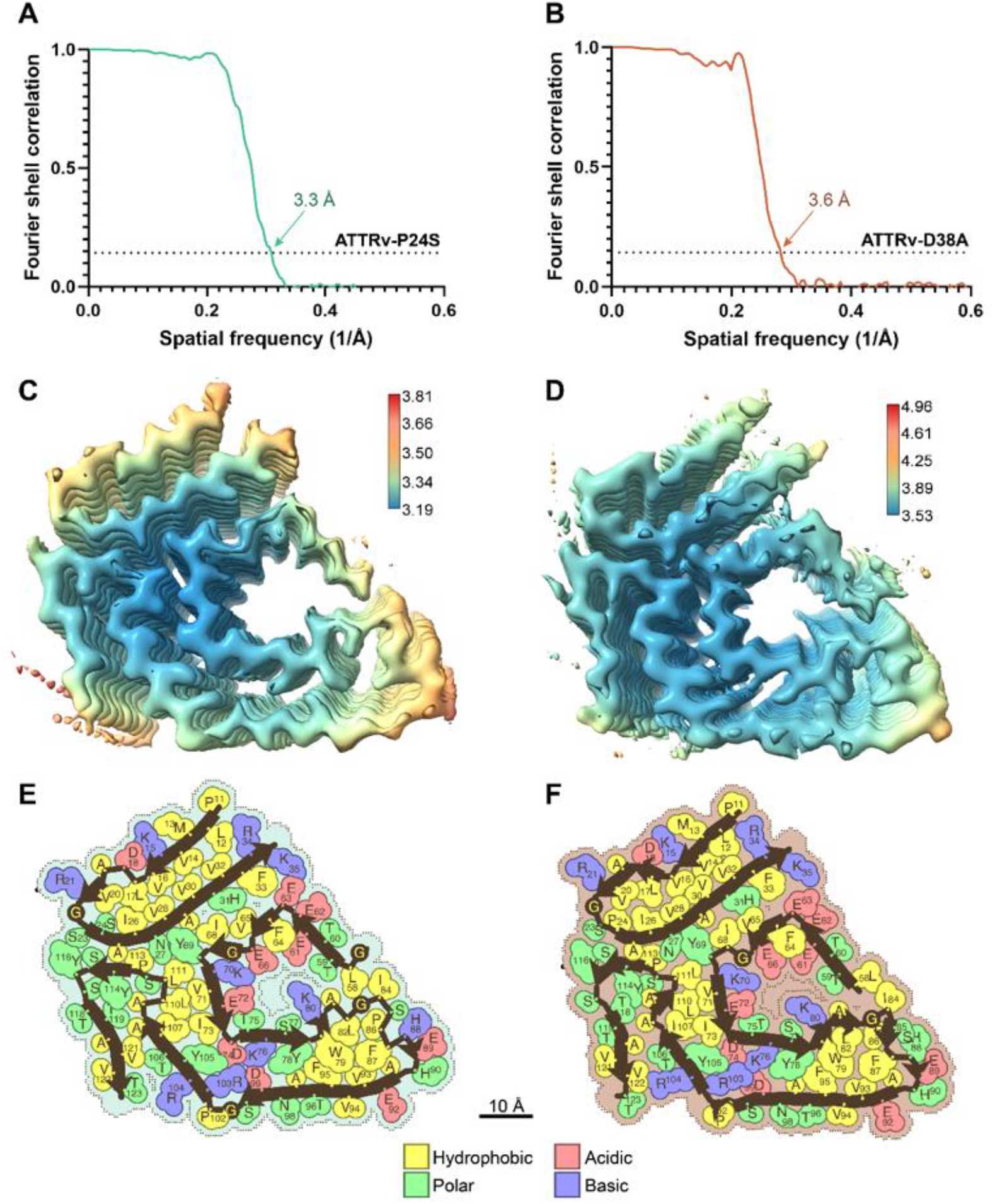
FSC curves and additional structural features. **(A and B)** Evaluation of the resolution of cryo-EM maps by Fourier shell correlation (FSC) curves of two independently refined half-maps from the ATTRv fibril structures of the two patients, ATTRv-P24S with resolution of 3.3 Å **(A)**, and ATTRv-D38A with resolution of 3.6 Å **(B). (C and D)** Local resolution maps obtained from ATTRv-P24S **(C)** and ATTRv-D38A **(C)** fibrils. Resolution scale in Angstroms; blue represents high resolution and red represents low resolution. **(E and F)** Schematic view of ATTRv-P24S **(E)** and ATTRv-D38A **(F)** fibrils showing residue composition. Residues are color coded by amino acid category, as labeled.

### Proteolytic and structural comparison of the ATTRv fibrils

The cryo-EM models enabled the mapping of non-tryptic proteolytic sites withing the ATTR structure **(Figure 5A, Supplementary Table 2)**. The three variants share common cleavage sites, while others have only been observed in a specific variant. Proteolysis was found not only in the solvent-exposed regions of the fibrils but also within the structured core. To explore whether these cleavage events might be related to local conformational perturbations or changes in stability, we aligned the fibril backbones and analyzed the solvation energy profiles of each variant.

**Figure 5.**
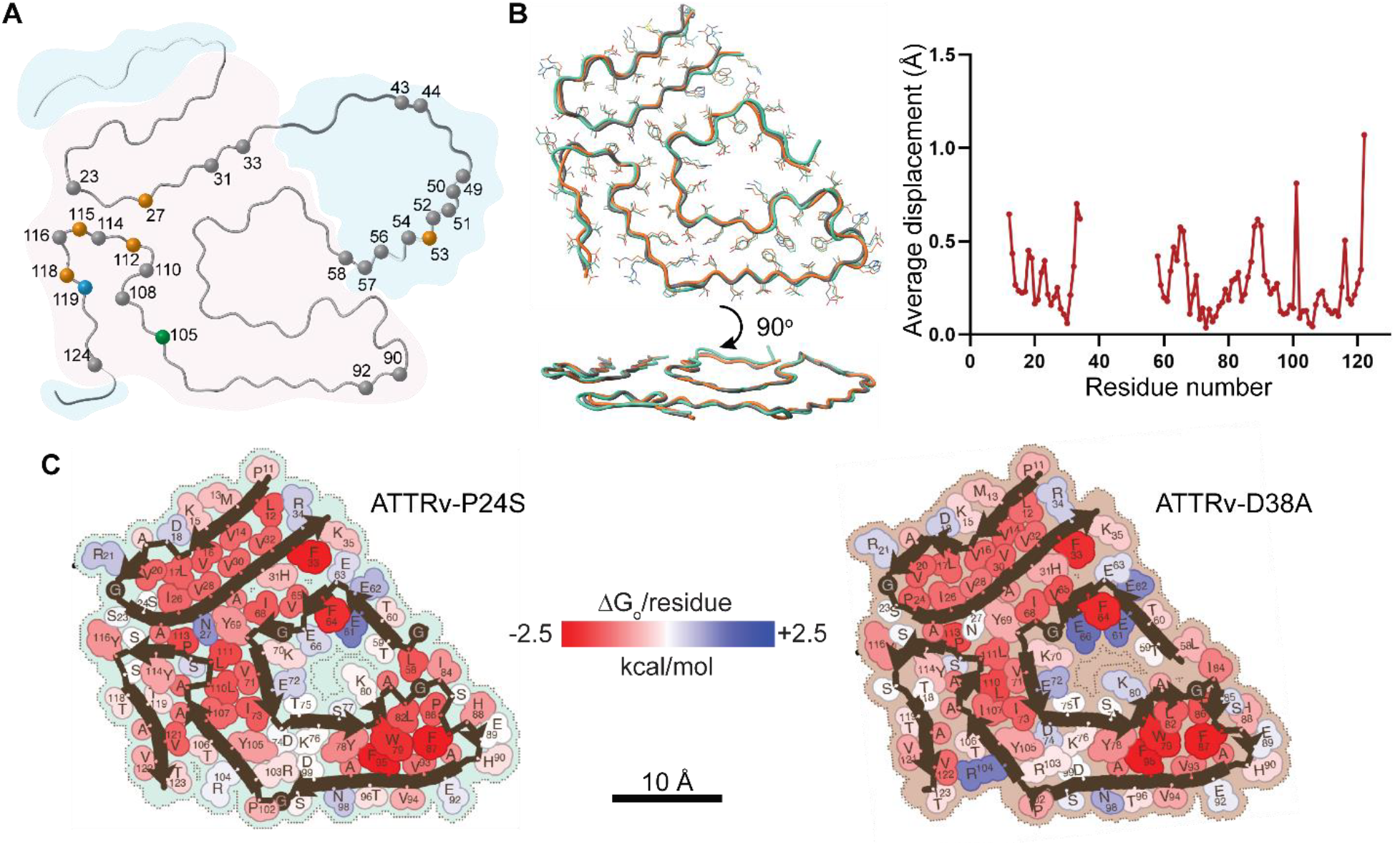
Proteolytic mapping, structural alignment, and solvation analysis. (A) Illustration summarizing the location of non-tryptic proteolytic sites within the resolved structures. Residues forming the closed-gate structure are highlighted with a pink shadow. Residues absent from the model but detected by mass spectrometry, presumably in an unfolded state, are shaded in blue. Common proteolytic sites shared across the three variants are marked in grey, while those unique to ATTRv fibrils carrying the P24S, D38A, or A25S mutations are indicated with green, orange, and blue dots, respectively. **(B)** Left, Structural alignment of ATTRv-P24S, ATTRv-D38A and ATTRwt (PDB 8e7d); top view and side view showing near identical overlapping conformation. Right, the average displacement value comparing fibril structures of ATTRv-P24S, ATTRv-D38A and five published ATTRwt structures. **(C)** Representation of solvation energies per residue estimated from ATTRv-P24S and ATTRv-D38A fibril structures. Residues are colored from favorable (red, -2.5 kcal/mol) to unfavorable stabilization energy (blue, 2.5 kcal/mol). Scale, 10 Å.

The Cα backbone structural alignment of ATTRv-P24S and ATTRv-D38A to an ATTRwt fibril structure (PDB 8e7d) revealed nearly identical structures in both top and side views **(Figure 6A)**. This high degree of similarity was confirmed by pairwise root-mean-square deviation (r.m.s.d.) comparisons with a consensus sequence derived from averaging r.m.s.d. of four ATTRwt structures (PDB codes 8gbr, 8e7d, 8e7h, 8ade), ATTRv-P24S and ATTRv-D38A, generated by GESAMT method.^24^ The r.m.s.d. values showed minimal displacements of 0.639 Å (ATTRv-P24S) and 0.586 Å (ATTRv-D38A) **(Supplementary Table 4)**. Residue-by-residue analysis further supports this structural similarity by revealing most displacements to be under 0.5 Å **(Figure 5B)**.

The estimated solvation energies of ATTRv-P24S, ATTRv-D38A, and ATTRwt are also comparable to previously described structures of ATTRv and ATTRwt fibrils. The fibril structure of ATTRv-P24S exhibited free energies per chain and per residue of -65.8 kcal/mol and -0.72 kcal/mol, respectively, while ATTRv-D38A had values of -60.6 kcal/mol and -0.67 kcal/mol **(Figure 6C, D)**. These stability profiles suggest fibril stabilities comparable to those previously published ATTRv and ATTRwt fibrils.^9, 17, 20, 21, 25^

## Discussion

The molecular basis of the phenotypic variability observed in patients with ATTRv amyloidosis remains elusive. We previously found that the neuropathy-associated mutations ATTRv-I84S and ATTRv-V122Δ drive the formation of polymorphic conformations.^9, 21^ Motivated by these findings, we set out to investigate a potential relationship between fibril polymorphism and neuropathic symptoms in ATTRv amyloidosis. We selected three patients carrying the mutations ATTRv-D38A, ATTRv-P24S, or ATTRv-A25S, all of whom presented with some form of neuropathy.^26-29^ Using cryo-EM, we determined the structures of their cardiac fibrils and evaluated their structural features **(Figures 2-6)**. Below, we discuss the implications of these results.

To date, all ATTR mutations studied have exhibited a single predominant fibril conformation in the heart, and only two mutations have been linked to additional conformations.^9, 16-21^ Roughly 19 cryo-EM structures of cardiac amyloid fibrils have been reported, and most fibrils extracted from cardiac tissue display a nearly identical single-protofilament architecture across nine genotypes and diverse patient demographics.^16-21, 25^ We refer to this prevalent architecture as the “closed-gate” conformation, distinguished by a polar channel running perpendicular to the fibril axis. The structures described here adopt the same fold, with no evidence of additional fibril conformations **(Figure 3 and 4)**.^9, 16-21^ So far, only ATTRv-I84S, which shows local variations in the polar channel,^9^ and ATTRv-V122Δ, which forms multi-protofilament fibrils^30^ deviate from this pattern. Whether these structural variations explain phenotypic differences remains an exciting open question.

Our study presents several limitations, each matched by meaningful opportunities for future investigation. The first limitation of our study is the limited number of patients and the intrinsic clinical diversity of ATTR amyloidosis. This inevitably complicates structure-phenotype analyses, but it also highlights the need for larger, systematic cohorts. Although all three patients studied here exhibited neuropathy, their presentations differed markedly **(Supplementary Table 1)**.^26-29^ The ATTRv-P24S patient presented with carpal tunnel syndrome, peripheral neuropathy, and cardiomyopathy.^26^ The ATTRv-A25S patient showed a unique clinical presentation with an acute onset neuropathy days after an influenza vaccination, progressing to severe symptomatology within 2 years.^27^ The case report describing this patient, which is the only one ever published for this mutation, also mentions that a sibling carrying the same mutation presented with dysesthesias in the lower extremities, electrocardiogram abnormalities, and 99mTc-scan consistent with cardiac amyloidosis, although she did not have nerve conduction abnormalities or symptoms of heart disease at the time of evaluation.^27^ Finally, the ATTRv-D38A patient presented with severe neuropathy, consistent with previous reports ^28, 29^. This patient underwent a liver transplantation, which was followed by rapidly progressive cardiac involvement, ultimately necessitating heart transplantation a few years after the first surgical intervention. Such variability must be considered when interpreting structural findings and underscores the value of expanding future studies to include broader clinical spectra.

A second limitation is that most mutations characterized so far occupy fibril regions that appear structurally tolerant. Unsurprisingly, variants such as ATTRv-D38A **(Figure 4C)** and ATTRv-V20I, G47E, T60A, and V122I ^31, 32^ do not perturb the core fold because they lie outside key stability pockets **(Figure 6C and D)**. Even ATTRv-V122Δ, while inducing multi-protofilament assembly, leaves the protofilament fold unchanged.^30^ Conversely, ATTRv-I84S—which disrupts a hydrophobic contact with Leu 58—produces subtle local rearrangements within the polar channel.^9^ However, mutations such as ATTRv-P24S and ATTRv-A25S insert polar residues into a hydrophobic interface yet still preserve the overall fold **(Figure 6C)**. One explanation is that multilayer contacts in the mature fibril buffer single-residue effects.^33^ Another is that abundant wild-type TTR within heterozygous fibrils masks localized perturbations. Both possibilities warrant further investigation.

A third limitation is that all patients examined to date are heterozygous, and mass - pectrometry confirms substantial wild-type content in their fibrils **(Supplementary Table 3)**.^17, 18, 20, 25^ It is possible that the structural changes driven by the single mutations ATTRv-D38A, ATTRv-P24S, or ATTRv-A25S described here or ATTRv-V30M described elsewhere^34, 35^ may not generate a sufficient disruption to overcome the structural influence of the wild-type conformation. Alternatively, the mutations could trigger localized structural changes that lack uniformity throughout the fibril. Consequently, these structures may represent primarily wild-type conformations. This possibility is particularly relevant when considering the intricacies of cryo-EM data processing, which relies on averaging. If the mutation introduces structural heterogeneity within the fibrils, such information is likely lost during the data-averaging process. Future studies of homozygous cases, though rare, will be critical for isolating mutation-driven effects.

Finally, we note that the present work interrogates fibrils purified exclusively from cardiac tissue. Although all three patients exhibited neuropathic involvement, amyloid in peripheral nerves or other organs was not examined and could, with a different local milieu, adopt alternative architectures. Thus, our conclusion that we observed no polymorphism linked to neuropathy cases so far applies only to fibrils taken from the heart. But we should also consider our earlier cryo-EM work on ATTRv-V30M, where fibrils taken separately from heart muscle and sural nerve shared the same closed-gate structure.^34^ This finding argues that organ-specific environments do not inevitably dictate different fibril folds. Even so, systematic multi-tissue analyses across a broader range of variants will be essential to determine whether the structural uniformity observed for V30M is the rule or an exception, thereby clarifying how (or if) tissue context shapes amyloid architecture in ATTRv amyloidosis.

Despite these challenges, our data reinforce a central observation: ATTRv cardiac fibrils adopt a remarkably conserved architecture across mutations and clinical presentations. Whether this homogeneity itself influences disease course, or whether subtler, localized polymorphisms play a role, remains to be clarified. Our work adds three new structures to the ATTR catalogue and emphasizes both the stability of the closed-gate fold and the promise of cryo-EM for disentangling genotype-phenotype links in amyloidosis, should there be any. Continued structural efforts, especially in homozygous or more clinically homogeneous cases, will help illuminate how specific fibril features contribute to the diverse manifestations of ATTRv amyloidosis.

## Acknowledgements

In memory of Dr. Merrill D. Benson, whose significant contributions have advanced our understanding of amyloid diseases and provided invaluable support to affected families for many years. We are grateful to the patients and families who generously donated tissues and to the University of Indiana for providing these materials. We appreciate the engaging discussions and insightful feedback from SBWIP and UTSW. We are thankful to the UTSW Cryo-Electron Microscopy Facility, UTSW Structural Biology Laboratory, and UTSW Electron Microscopy Core Facility, as well as the national cryo-EM facilities at Stanford-SLAC (project CA60) and PNCC (project 51267) for their instrumental and technical support, and data collection efforts. We also extend our gratitude to the UTSW Proteomics core for their technical support in the proteomics experiments.

## Funding

American Heart Association, Career Development Award 847236, L.S.

National Institutes of Health, National Heart, Lung, and Blood Institute, New Innovator Award DP2-HL163810, L.S.

Welch Foundation, Research Award I-2121-20220331, L.S.

UTSW Endowment, Distinguished Researcher Award from President’s Research Council and start-up funds, L.S.

Cryo-EM research was partially supported by the following grants:

National Institutes of Health grant U24GM129547, Department of Energy Office of Science User Facility sponsored by the Office of Biological and Environmental Research

Department of Energy, Laboratory Directed Research and Development program at SLAC National Accelerator Laboratory, under contract DE-AC02-76SF00515

NIH Common Fund Transformative High Resolution Cryo-Electron Microscopy program (U24 GM129539)

The Cryo-Electron Microscopy Facility and the Structural Biology Laboratory at UTSW are supported by a grant from the Cancer Prevention & Research Institute of Texas (RP170644).

The Electron Microscopy Core Facility at UTSW is supported by the National Institutes of Health (NIH) (1S10OD021685-01A1 and 1S10OD020103-01).

Part of the computational resources were provided by the BioHPC supercomputing facility located in the Lyda Hill Department of Bioinformatics at UTSW. URL: https://portal.biohpc.swmed.edu.

## Author contributions

Conceptualization: B.N., S.A., M.C.F.R., L.S.

Methodology: B.N., S.A., M.C.F.R.

Investigation: B.N., S.A., M.C.F.R., M.P., P.S., Y.A., R.P., A.L., F.C., B.K.B., D.E., L.S.

Visualization: B.N., S.A., M.C.F.R., L.S.

Funding acquisition: L.S.

Project administration L.S.

Supervision: B.N., L.S.

Writing – original draft: V.S., L.S.

Writing – review & editing: B.N., S.A., M.C.F.R., L.S.

## Declaration of interests

L.S. reports research funding from NHLBI, Welch Foundation, UTSW, and AstraZeneca. L.S. also reports advisory board, speaker, and consulting fees from Alexion, Pfizer, Attralus, Intellia, and AmyGo. B.A.N. also reports advisory board, speaker, and consulting fees from Amygo. L.S, R.P and B.A.N are the co-founders of AmyGo solutions. D.S.E. is chair of the scientific advisory board of and an equity holder in ADRx.

## Data and Materials Availability

Structural data have been deposited into the Worldwide Protein Data Bank (wwPDB) and the Electron Microscopy Data Bank (EMDB) with the following EMD accession codes: EMD-26692 (ATTRv-P24S-), EMD-49581 (ATTRv-A25S), EMD-49578 (ATTRv-D38A) and PDB accession codes: 8e7i (ATTRv-P24S) and 9nnn (ATTRvD-38A). All data generated or analyzed during this study that support the findings are available within this published article and its supplementary data files. Graphed data is provided in the source data file. Cardiac specimens were obtained from the laboratory of the late Dr. Merrill D. Benson at Indiana University or from UCLA. These specimens are under a material transfer agreement with the universities and cannot be distributed freely.

## Methods

### Patients and tissue material

We obtained fresh frozen or lyophilized cardiac tissues from three ATTRv amyloidosis patients. All samples were postmortem. The details of each tissue sample included in the study are listed in Supplementary Table 1. Specimens from the left ventricle of either explanted or autopsied hearts were obtained from the laboratory of the late Dr. Merrill D. Benson at the University of Indiana and Mario Deng, UCLA. The Office of the Human Research Protection Program granted exemption from Internal Review Board review because all specimens were anonymized.

### Extraction of amyloid fibrils from human cardiac tissue

*Ex-vivo* preparations of amyloid fibrils were obtained from the fresh-frozen or lyophilized human tissue as described earlier. ^36^ Briefly, ∼200 mg of frozen/lyophilized cardiac tissue per patient was thawed at room temperature and cut into small pieces with a scalpel. The minced tissue was resuspended into 1 mL Tris-calcium buffer (20 mM Tris, 138 mM NaCl, 2 mM CaCl_2_, 0.1% NaN_3_, pH 8.0) and centrifuged for 5 min at 3100 × g and 4 °C. The pellet was washed in Tris-calcium buffer four additional times. After the washing, the pellet was resuspended in 1 mL of 5 mg mL^-1^ collagenase solution (collagenase stock prepared in Tris-calcium buffer) and incubated overnight at 37 °C, shaking at 400 rpm. The resuspension was centrifuged for 30 min at 3100 × g and 4 °C and the pellet was resuspended in 1 mL Tris–ethylenediaminetetraacetic acid (EDTA) buffer (20 mM Tris, 140 mM NaCl, 10 mM EDTA, 0.1% NaN_3_, pH 8.0). The suspension was centrifuged for 5 min at 3100 × g and 4 °C, and the washing step with Tris–EDTA was repeated nine additional times. All the supernatants were collected for further analysis, when needed. After the washing, the pellet was resuspended in 200 μL ice-cold water supplemented with 5 mM EDTA and centrifuged for 5 min at 3100 × g and 4 °C. This step released the amyloid fibrils from the pellet, which were collected in the supernatant. This extraction step was repeated three additional times. The material from the various patients was handled and analyzed separately.

### Negative-stained transmission electron microscopy

Amyloid fibril extraction was confirmed by transmission electron microscopy as described[ref]. Briefly, a 3 μL sample was spotted onto a freshly glow-discharged carbon film 300 mesh copper grid (Electron Microscopy Sciences), incubated for 2 min, and gently blotted onto a filter paper to remove the solution. The grid was negatively stained with 3 µL of 2% uranyl acetate for 2 min and gently blotted to remove the solution. Another 3 μL uranyl acetate was applied onto the grid and immediately removed. An FEI Tecnai 12 electron microscope at an accelerating voltage of 120 kV was used to examine the specimens.

### Western blotting of extracted ATTR fibrils

To identify the fibril type, western blotting was performed on the extracted fibrils. In summary, 0.5 µg of fibrils were dissolved in a tricine SDS sample buffer and heated to 85°C for 2 minutes. Type B fibrils were used as control. The samples were then applied to a Novex™ 16% tris-tricine gel system with a Tricine SDS running buffer. The fibril type (type A vs Type B) was determined by transferring the gel to a 0.2 µm nitrocellulose membrane, which was then probed with a primary antibody (diluted 1:1000) specific to the C-terminal region of the wild-type transthyretin sequence (GenScript). A secondary antibody, horseradish peroxidase-conjugated goat anti-rabbit IgG (Invitrogen, diluted 1:1000), was used for detection. Transthyretin content was visualized using Promega Chemiluminescent Substrate, as per the manufacturer’s guidelines.

### Mass Spectrometry (MS) sample preparation, data acquisition and analysis

For tryptic MS analysis, 0.5 µg of extracted ATTR fibrils were dissolved in a tricine SDS sample buffer, boiled for 2 minutes at 85 °C, and run on a Novex™ 16% tris-tricine gel system using a Tricine SDS running buffer. Gel was stained with Coomassie dye, destained and ATTR smear was cut from the gel. Sample was sent for MS analysis. Samples were digested overnight with trypsin (Pierce) following reduction and alkylation with DTT and iodoacetamide (Sigma–Aldrich). The samples then underwent solid-phase extraction cleanup with an Oasis HLB plate (Waters) and the resulting samples were injected onto an Q Exactive HF mass spectrometer coupled to an Ultimate 3000 RSLC-Nano liquid chromatography system. Samples were injected onto a 75 um i.d., 15-cm long EasySpray column (Thermo) and eluted with a gradient from 0-28% buffer B over 90 min. Buffer A contained 2% (v/v) ACN and 0.1% formic acid in water, and buffer B contained 80% (v/v) ACN, 10% (v/v) trifluoroethanol, and 0.1% formic acid in water. The mass spectrometer operated in positive ion mode with a source voltage of 2.5 kV and an ion transfer tube temperature of 300 °C. MS scans were acquired at 120,000 resolution in the Orbitrap and up to 20 MS/MS spectra were obtained in the ion trap for each full spectrum acquired using higher-energy collisional dissociation (HCD) for ions with charges 2-8. Dynamic exclusion was set for 20 s after an ion was selected for fragmentation.

Raw MS data files were analyzed using Proteome Discoverer v3.0 SP1 (Thermo), with peptide identification performed using a semitryptic search with Sequest HT against the human reviewed protein database from UniProt. Fragment and precursor tolerances of 10 ppm and 0.02 Da were specified, and three missed cleavages were allowed. Carbamidomethylation of Cys was set as a fixed modification, with oxidation of Met set as a variable modification. The false-discovery rate (FDR) cutoff was 1% for all peptides. The mass spectrometry proteomics data have been deposited to MassIVE (a member of ProteomeXchange) with accession number MSV000098698.

### Cryo-EM sample preparation, data collection, and processing

Freshly extracted fibril samples were applied to glow-discharged Quantifoil R 1.2/1.3, 300 mesh, Cu grids, blotted with filter paper to remove excess sample, and plunged frozen into liquid ethane using a Vitrobot Mark IV (FEI). Cryo-EM samples were screened on either the Talos Arctica or Glacios at the Cryo-Electron Microscopy Facility (CEMF) at The University of Texas Southwestern Medical Center (UTSW), and the final datasets were collected on a 300 kV Titan Krios microscope (FEI) at the Stanford-SLAC Cryo-EM Center (S^2^C^2^) (Supplementary Table 5). Pixel size, frame rate, dose rate, final dose, and number of micrographs per sample are detailed in Table S4. Automated data collection was performed by SerialEM software package^37^ and Thermo Scientific Smart EPU software. The raw movie frames were gain-corrected, aligned, motion-corrected and dose-weighted using RELION’s own implemented motion correction program.^38^ Contrast transfer function (CTF) estimation was performed using CTFFIND 4.1.^39^ All steps of helical reconstruction, three-dimensional (3D) refinement, and post-process were carried out using RELION 3.1 and RELION 4.0. ^40, 41^ Filaments for ATTRv-P24S and ATTRv-D38A were manually picked using EMAN2 e2helixboxer.py, while filaments for ATTRv-A25S were manually picked using RELION4.0.^42^ Particles were extracted using a box size of 1024 and 256 pixels with an inter-box distance of 10% of the box length. 2D classification of 1024-pixel particles was used to estimate the helical parameters. 2D classifications of 256-pixel particles were used to select suitable particles for further processing. Fibril helix is assumed to be left-handed. We used an elongated Gaussian blob as an initial reference for 3D classifications of ATTRv. For ATTRv, we used a RELION built-in script, *relion_helix_inimodel2d*, to generate an incomplete initial model and low-pass filtered the model to 50 Å. We performed 3D classifications with an average of ∼30k to 40k particles per class to separate filament types. Particles potentially leading to the best reconstructed map were chosen for 3D auto-refinements. CTF refinements and Bayesian polishing were performed to obtain higher resolution. Final maps were post-processed using the recommended standard procedures in RELION. The final subset of selected particles was used for high-resolution gold-standard refinement as described previously.^43^ The final overall resolution estimate was evaluated based on the FSC at 0.143 threshold between two independently refined half-maps (Supplementary Figure 3).^44^

### Model building

The refined maps were further sharpened using phenix.auto_sharpen at the resolution cutoff.^45^ Our previously published model of ATTRwt (PDB 8e7d) was used as the template to build the model of ATTRv-P24S, which in turn, was used as a template to build the other ATTRv models. Residue modification, rigid body fit zone, and real space refine zone were performed to obtain the resulting models using COOT.^46^ All the statistics are summarized in Supplementary Table 5.

### Stabilization energy calculation

The stabilization energy per residue was calculated by the sum of the products of the area buried for each atom and the corresponding atomic solvation parameters.^47, 48^ The overall energy was calculated by the sum of energies of all residues, and assorted colors were assigned to each residue, instead of each atom, in the solvation energy map.

### Figure Panels

All figure panels were created with ChimeraX 1.8 and Adobe illustrator.

## References

1. Y. Ando, T. Coelho, J. L. Berk, M. W. Cruz, B.-G. Ericzon, S.-i. Ikeda, W. D. Lewis, L. Obici, V. Planté-Bordeneuve, C. Rapezzi, G. Said and F. Salvi, Orphanet Journal of Rare Diseases, 2013, 8, 31.

2. A. Dispenzieri, T. Coelho, I. Conceição, M. Waddington-Cruz, J. Wixner, A. V. Kristen, C. Rapezzi, V. Planté-Bordeneuve, J. Gonzalez-Moreno, M. S. Maurer, M. Grogan, D. Chapman, L. Amass, P. G. Pavia, I. Tarnev, J. G. Costello, M. A. G. D. Briseno, H. Schmidt, B. Drachman, F. A. Barroso, T. Yamashita, O. Lairez, Y. Sekijima, G. Vita, E.-S. Jeon, M. Hanna, D. Slosky, M. Luigetti, S. LoRusso, F. M. Beamud, D. Adams, H. Moelgaard, R. Press, C. L. Cirami, H. Nienhuis, J. M. C. Plana, J. Inamo, D. Jacoby, M. Emdin, D. Quan, S. Hummel, R. Witteles, A. Dori, S. Shah, D. Lenihan, O. Azevedo, S. Murali, S. Zivkovic, S. C. Low, J. Nativi-Nicolau, N. Fine, J. Tallaj, C. Tschoepe, R. F. Torrón, M. Polydefkis, G. Merlini, S. Badelita, S. Gottlieb, J. Tauras, E. B. Correia, H. Ventura, B. Gess, F. Darstein, J. Oh, T. Marburger, J. Van Cleemput, V. L. Salutto, Y. Parman, C.-C. Chao, N. Sarswat, C. Mueller, D. Steidley, J. Ralph, A. Warner, W. Cotts, J. Hoffman, M. Rugiero, S. Misawa, J. L. M. Blanco, L. G. Davila, M. Sadeh, J. Luo, T. Kyriakides, A. Wang, H. Kaufmann, S. Zivkovic and T. i. the, Orphanet Journal of Rare Diseases, 2022, 17, 236.

3. E. González-López, M. Gallego-Delgado, G. Guzzo-Merello, F. J. de Haro-del Moral, M. Cobo-Marcos, C. Robles, B. Bornstein, C. Salas, E. Lara-Pezzi, L. Alonso-Pulpon and P. Garcia-Pavia, European Heart Journal, 2015, 36, 2585–2594.

4. A. Carroll, P. J. Dyck, M. d. Carvalho, M. Kennerson, M. M. Reilly, M. C. Kiernan and S. Vucic, Journal of Neurology, Neurosurgery & Psychiatry, 2022, 93, 668–678.

5. S. Zampino, F. H. Sheikh, J. Vaishnav, D. Judge, B. Pan, A. Daniel, E. Brown, G. Ebenezer and M. Polydefkis, Neurology, 2023, 100, e2036–e2044.

6. Z. L. Almeida, D. C. Vaz and R. M. M. Brito, Crit Rev Clin Lab Sci, 2024, 61, 616–640.

7. E. Carvalho, A. Dias, T. Coelho, A. Sousa, M. Alves-Ferreira, M. Santos and C. Lemos, J Neurol, 2024, 271, 5746–5761.

8. A. Porcari, M. Fontana and J. D. Gillmore, Cardiovasc Res, 2023, 118, 3517–3535.

9. B. A. Nguyen, V. Singh, S. Afrin, A. Yakubovska, L. Wang, Y. Ahmed, R. Pedretti, M. D. C. Fernandez-Ramirez, P. Singh, M. Pękała, L. O. Cabrera Hernandez, S. Kumar, A. Lemoff, R. Gonzalez-Prieto, M. R. Sawaya, D. S. Eisenberg, M. D. Benson and L. Saelices, Nat Commun, 2024, 15, 581.

10. A. W. P. Fitzpatrick, B. Falcon, S. He, A. G. Murzin, G. Murshudov, H. J. Garringer, R. A. Crowther, B. Ghetti, M. Goedert and S. H. W. Scheres, Nature, 2017, 547, 185–190.

11. S. H. W. Scheres, B. Ryskeldi-Falcon and M. Goedert, Nature (London), 2023, 621, 701–710.

12. S. H. Scheres, W. Zhang, B. Falcon and M. Goedert, Curr Opin Struct Biol, 2020, 64, 17–25.

13. Y. Yang, D. Arseni, W. Zhang, M. Huang, S. Lövestam, M. Schweighauser, A. Kotecha, A. G. Murzin, S. Y. Peak-Chew, J. Macdonald, I. Lavenir, H. J. Garringer, E. Gelpi, K. L. Newell, G. G. Kovacs, R. Vidal, B. Ghetti, B. Ryskeldi-Falcon, S. H. W. Scheres and M. Goedert, Science, 2022, 375, 167–172.

14. Y. Yang, H. J. Garringer, Y. Shi, S. Lövestam, S. Peak-Chew, X. Zhang, A. Kotecha, M. Bacioglu, A. Koto, M. Takao, M. G. Spillantini, B. Ghetti, R. Vidal, A. G. Murzin, S. H. W. Scheres and M. Goedert, Acta Neuropathol, 2023, 145, 561–572.

15. Y. Yang, Y. Shi, M. Schweighauser, X. Zhang, A. Kotecha, A. G. Murzin, H. J. Garringer, P. W. Cullinane, Y. Saito, T. Foroud, T. T. Warner, K. Hasegawa, R. Vidal, S. Murayama, T. Revesz, B. Ghetti, M. Hasegawa, T. Lashley, S. H. W. Scheres and M. Goedert, Nature, 2022, 610, 791–795.

16. M. Steinebrei, J. Gottwald, J. Baur, C. Röcken, U. Hegenbart, S. Schönland and M. Schmidt, Nat Commun, 2022, 13, 6398.

17. B. A. Nguyen, V. Singh, S. Afrin, P. Singh, M. Pekala, Y. Ahmed, R. Pedretti, J. Canepa, A. Lemoff, B. Kluve-Beckerman, P. M. Wydorski, F. Chhapra and L. Saelices, Communications Biology, 2024, 7, 905.

18. M. Schmidt, S. Wiese, V. Adak, J. Engler, S. Agarwal, G. Fritz, P. Westermark, M. Zacharias and M. Fändrich, Nat Commun, 2019, 10, 5008.

19. M. Steinebrei, J. Baur, A. Pradhan, N. Kupfer, S. Wiese, U. Hegenbart, S. O. Schönland, M. Schmidt and M. Fändrich, Nat Commun, 2023, 14, 7623.

20. M. D. C. Fernandez-Ramirez, B. A. Nguyen, V. Singh, S. Afrin, B. Evers, P. Basset, L. Wang, M. Pękała, Y. Ahmed, P. Singh, J. Canepa, A. Wosztyl, Y. Li and L. Saelices, bioRxiv, 2024, DOI: 10.1101/2024.05.14.594218.

21. Y. Ahmed, B. A. Nguyen, S. Afrin, V. Singh, B. Evers, P. Singh, R. Pedretti, L. Wang, P. Bassett, M. d. C. Fernandez-Ramirez, M. Pekala, B. Kluve-Beckerman and L. Saelices, bioRxiv, 2024, DOI: 10.1101/2024.05.09.593396, 2024.2005.2009.593396.

22. E. Ihse, A. Ybo, O. Suhr, P. Lindqvist, C. Backman and P. Westermark, J Pathol, 2008, 216, 253–261.

23. E. Ihse, C. Rapezzi, G. Merlini, M. Benson, Y. Ando, O. Suhr, S. Ikeda, F. Lavatelli, L. Obici, C. Quarta, O. Leone, H. Jono, M. Ueda, M. Lorenzini, J. Liepnieks, T. Ohshima, M. Tasaki, T. Yamashita and P. Westermark, Amyloid, 2013, 20, 142–150.

24. E. Krissinel, J Mol Biochem, 2012, 1, 76–85.

25. B. A. Nguyen, S. Afrin, A. Yakubovska, V. Singh, J. V. Alicea, P. Kunach, P. Singh, M. Pekala, Y. Ahmed, M. d. C. Fernandez-Ramirez, L. O. C. Hernandez, R. Pedretti, P. Bassett, L. Wang, A. Lemoff, L. Villalon, B. Kluve-Beckerman and L. Saelices, bioRxiv, 2024, DOI: 10.1101/2024.05.14.594028, 2024.2005.2014.594028.

26. T. Uemichi, M. A. Gertz and M. D. Benson, J Med Genet, 1995, 32, 279–281.

27. M. Yazaki, T. Yamashita, J. C. Kincaid, J. R. Scott, R. G. Auger, P. J. Dyck and M. D. Benson, Muscle Nerve, 2002, 25, 244–250.

28. H. J. Cho, J. Y. Yoon, M. H. Bae, J. H. Lee, D. H. Yang, H. S. Park, Y. Cho, S. C. Chae and J. E. Jun, J Cardiovasc Ultrasound, 2012, 20, 209–212.

29. M. Kishikawa, T. Nakanishi, A. Miyazaki, A. Shimizu, H. Kusaka, M. Fukui and T. Nishiue, Amyloid, 1999, 6, 278–281.

30. Y. Ahmed, B. A. Nguyen, S. Afrin, V. Singh, B. Evers, P. Singh, R. Pedretti, L. Wang, P. Bassett, M. Del Carmen Fernandez-Ramirez, M. Pekala, B. Kluve-Beckerman and L. Saelices, bioRxiv, 2024, DOI: 10.1101/2024.05.09.593396.

31. S. Schönland, M. Schmidt and M. Fändrich, Nature Communications, 2023.

32. M. d. C. Fernandez-Ramirez, B. A. Nguyen, V. Singh, S. Afrin, B. M. Evers, P. Bassett, L. Wang, M. Pekala, Y. Ahmed, P. Singh, J. Canepa, A. Wosztyl, Y. Li and L. Saelices, bioRxiv, 2024, DOI: 10.1101/2024.05.14.594218, 2024.2005.2014.594218.

33. M. Sawaya, M. Hughes, J. Rodriguez, R. Riek and D. Eisenberg, Cell, 2021, 184, 4857–4873.

34. B. A. Nguyen, S. Afrin, A. Yakubovska, V. Singh, J. Vaquer-Alicea, P. Kunach, P. Singh, M. Pekala, Y. Ahmed, M. d. C. Fernandez-Ramirez, L. C. Hernandez, R. Pedretti, P. Bassett, L. Wang, A. Lemoff, L. Villalon, B. Kluve-Beckerman and L. S. Gomez, Structure, 2024, DOI: 10.1016/j.str.2024.09.021, in press.

35. M. Schmidt, S. Wiese, V. Adak, J. Engler, S. Agarwal, G. Fritz, P. Westermark, M. Zacharias and M. Fändrich, Nat Commun, 2019, 10, 5008.

36. B. A. Nguyen, V. Singh, S. Afrin, A. Yakubovska, L. Wang, Y. Ahmed, R. Pedretti, M. d. C. Fernandez-Ramirez, P. Singh, M. Pękała, L. O. Cabrera Hernandez, S. Kumar, A. Lemoff, R. Gonzalez-Prieto, M. R. Sawaya, D. S. Eisenberg, M. D. Benson and L. Saelices, Nature Communications, 2024, 15, 581.

37. D. N. Mastronarde, J Struct Biol, 2005, 152, 36–51.

38. J. Zivanov, T. Nakane and S. H. W. Scheres, IUCrJ, 2019, 6, 5–17.

39. A. Rohou and N. Grigorieff, J Struct Biol, 2015, 192, 216–221.

40. D. Kimanius, L. Dong, G. Sharov, T. Nakane and S. H. W. Scheres, Biochem J, 2021, 478, 4169–4185.

41. J. Zivanov, T. Nakane, B. O. Forsberg, D. Kimanius, W. J. Hagen, E. Lindahl and S. H. Scheres, Elife, 2018, 7.

42. J. M. Bell, M. Chen, T. Durmaz, A. C. Fluty and S. J. Ludtke, J Struct Biol, 2018, 204, 283–290.

43. S. H. W. Scheres, Journal of Structural Biology, 2012, 180, 519–530.

44. S. Chen, G. McMullan, A. R. Faruqi, G. N. Murshudov, J. M. Short, S. H. Scheres and R. Henderson, Ultramicroscopy, 2013, 135, 24–35.

45. T. C. Terwilliger, O. V. Sobolev, P. V. Afonine and P. D. Adams, Acta Crystallogr D Struct Biol, 2018, 74, 545–559.

46. P. Emsley, B. Lohkamp, W. G. Scott and K. Cowtan, Acta Crystallogr D Biol Crystallogr, 2010, 66, 486–501.

47. D. Eisenberg and A. D. McLachlan, Nature, 1986, 319, 199–203.

48. M. R. Sawaya, M. P. Hughes, J. A. Rodriguez, R. Riek and D. S. Eisenberg, Cell, 2021, 184, 4857–4873.

