## Supplemental Tables for "A Shared Amyloid Architecture in Cardiac Fibrils from Three Neuropathy-Associated ATTR Variants"

**Affiliations:**

**Supplementary Tables:**

**Supplementary Table 1.** List of ATTRv cardiac samples included in the study.

| Genotype | Origin | Sex | Age at death/ Autopsy | Neuropathy signs | Remarks |
|---|---|---|---|---|---|
| ATTRv-P24S | Transplant | Male | 69 | Yes | Cardiomyopathy, neuropathy, GI distress, carpal tunnel syndrome polyneuropathy[1] |
| ATTRv-A25S | Autopsy (Lyophilized) | Female | 54 | Yes | Polyneuropathy after receiving Flu vaccine at age 52. Severe and rapid progression resulting in death within 2 years. GI distress.[2] Symptoms in family members with same genotype: cardiomyopathy, longstanding toe numbness, carpal tunnel syndrome. |
| ATTRv-D38A | Transplant (Lyophilized) | Female | 59 | Unknown | Common symptoms in other patients: polyneuropathy, cardiomyopathy, autonomic function abnormalities [3, 4]. |

**Supplementary Table 2.** Complete list of tryptic and semi-tryptic transthyretin peptides [P02766] identified by MS analysis, with 1% False Discovery Rate (FDR) filtering. Mutations are highlighted in red.

| ATTRv-P24S | ATTRv-A25S | ATTRv-D38A |
|---|---|---|
| [R].GSPAINVAVH.[V] | [R].GSPAINVAVH.[V] | [R].KAADDTWEP.[F] |
| [R].GSPAINVAVHVFR.[K] | [R].GSPAINVAVHVF.[R] | [R].KAADDTWEPFASGK.[T] |
| [R].GSSAINVAVH.[V] | [R].GSPAINVAVHVFR.[K] | [K].AADDTWEP.[F] |
| [R].GSSAINVAVHVFR.[K] | [R].GSPSINVAVH.[V] | [K].AADDTWEPFASGK.[T] |
| [K].AADDTWEP.[F] | [R].GSPSINVAVHVFR.[K] | [R].KAAADTWEPF.[A] |
| [K].AADDTWEPF.[A] | [S].PSINVAVHVFR.[K] | [K].AAADTWEP.[F] |
| [K].AADDTWEPFASGK.[T] | [K].AADDTWEPF.[A] | [K].AAADTWEPFASGK.[T] |
| [K].ALGISPFHEHAEVVFTANDSGPR.[R] | [K].AADDTWEPFASGK.[T] | [H].AEVVFTANDSGPR.[R] |
| [A].ALLSPYSYSTTAVVTNPK.[E] | [H].AEVVFTANDSGPR.[R] | [K].ALGISPFHEHAEVVFTANDSGPR.[R] |
| [A].ALLSPYSYSTTAVVTNPKE.[-] | [K].ALGISPFHEHAEVVFTANDSGPR.[R] | [A].ALLSPYSYSTTAVVTNPKE.[-] |
| [H].GLTTEEEFVEGIYK.[V] | [A].ALLSPYSYSTTAVVTNPK.[E] | [G].ELHGLTTEEEFVEGIYK.[V] |
| [R].KAADDTWEP.[F] | [A].ALLSPYSYSTTAVVTNPKE.[-] | [S].ESGELHGLTTEEEFVEGIYK.[V] |
| [R].KAADDTWEPF.[A] | [S].ESGELHGLTTEEEFVEGIYK.[V] | [S].GELHGLTTEEEFVEGIYK.[V] |
| [R].KAADDTWEPFASGK.[T] | [S].GELHGLTTEEEFVEGIYK.[V] | [S].GELHGLTTEEEFVEGIYKVEIDTK.[S] |
| [L].LSPYSYSTTAVVTNPKE.[-] | [H].GLTTEEEFVEGIYK.[V] | [H].GLTTEEEFVEGIYK.[V] |
| [G].LTTEEEFVEGIYK.[V] | [R].KAADDTWEPFASGK.[T] | [H].GLTTEEEFVEGIYKVEIDTK.[S] |
| [R].RYTIAALLSPY.[S] | [E].LHGLTTEEEFVEGIYK.[V] | [R].GSPAINVAVH.[V] |
| [R].RYTIAALLSPYSY.[S] | [L].LSPYSYSTTAVVTNPK.[E] | [R].GSPAINVAVHVF.[R] |
| [R].RYTIAALLSPYSYSTTAVVTN.[P] | [L].LSPYSYSTTAVVTNPKE.[-] | [R].GSPAINVAVHVFR.[K] |
| [R].RYTIAALLSPYSYSTTAVVTNPK.[E] | [G].LTTEEEFVEGIYKVEIDTK.[S] | [E].LHGLTTEEEFVEGIYK.[V] |
| [R].RYTIAALLSPYSYSTTAVVTNPKE.[-] | [R].RYTIAALLSPYSY.[S] | [L].LSPYSYSTTAVVTNPKE.[-] |
| [T].SESGELHGLTTEEEFVEGIYK.[V] | [R].RYTIAALLSPYSYSTT.[A] | [G].LTTEEEFVEGIYK.[V] |
| [E].SGELHGLTTEEEFVEGIYK.[V] | [R].RYTIAALLSPYSYSTTAVVTN.[P] | [G].LTTEEEFVEGIYKVEIDTK.[S] |
| [Y].STTAVVTNPK.[E] | [R].RYTIAALLSPYSYSTTAVVTNPK.[E] | [S].PAINVAVHVFR.[K] |
| [Y].STTAVVTNPKE.[-] | [R].RYTIAALLSPYSYSTTAVVTNPKE.[-] | [S].PYSYSTTAVVTNPKE.[-] |
| [Y].SYSTTAVVTNPK.[E] | [T].SESGELHGLTTEEEFVEGIYK.[V] | [R].RYTIAALLSPY.[S] |
| [Y].SYSTTAVVTNPKE.[-] | [E].SGELHGLTTEEEFVEGIYK.[V] | [R].RYTIAALLSPYSY.[S] |
| [Y].TIAALLSPYSYSTTAVVTNPKE.[-] | [Y].STTAVVTNPK.[E] | [R].RYTIAALLSPYSYST.[T] |
| [K].TSESGELHGLTTEEEFVEGIYK.[V] | [Y].STTAVVTNPKE.[-] | [R].RYTIAALLSPYSYSTTAVVTN.[P] |
| [K].TSESGELHGLTTEEEFVEGIYKVEIDTK.[S] | [Y].SYSTTAVVTNPK.[E] | [R].RYTIAALLSPYSYSTTAVVTNPK.[E] |
| [L].TTEEEFVEGIYK.[V] | [Y].SYSTTAVVTNPKE.[-] | [R].RYTIAALLSPYSYSTTAVVTNPKE.[-] |
| [R].YTIAALLSPYSY.[S] | [K].TSESGELHGLTTEEEFVEGIYK.[V] | [T].SESGELHGLTTEEEFVEGIYK.[V] |
| [R].YTIAALLSPYSYSTTAVVTN.[P] | [K].TSESGELHGLTTEEEFVEGIYKVEIDTK.[S] | [T].SESGELHGLTTEEEFVEGIYKVEIDTK.[S] |
| [R].YTIAALLSPYSYSTTAVVTNPK.[E] | [L].TTEEEFVEGIYK.[V] | [E].SGELHGLTTEEEFVEGIYK.[V] |
| [R].YTIAALLSPYSYSTTAVVTNPKE.[-] | [L].TTEEEFVEGIYKVEIDTK.[S] | [E].SGELHGLTTEEEFVEGIYKVEIDTK.[S] |
| | [E].VVFTANDSGPR.[R] | [Y].STTAVVTNPKE.[-] |
| | [R].YTIAALLSPYSY.[S] | [Y].SYSTTAVVTNPK.[E] |

| | | |
|---|---|---|
| | [R].YTIAALLSPYSYSTTAVVTN.[P] | [Y].SYSTTAVVTNPKE.[-] |
| | [R].YTIAALLSPYSYSTTAVVTNPK.[E] | [T].TAVVTNPKE.[-] |
| | [R].YTIAALLSPYSYSTTAVVTNPKE.[-] | [K].TSESGELHGLTTEEEFVEGIYK.[V] |
| | | [K].TSESGELHGLTTEEEFVEGIYKVEIDTK.[S] |
| | | [L].TTEEEFVEGIYK.[V] |
| | | [L].TTEEEFVEGIYKVEIDTK.[S] |
| | | [N].VAVHVFR.[K] |
| | | [E].VVFTANDSGPR.[R] |
| | | [R].YTIAALLSPY.[S] |
| | | [R].YTIAALLSPYSYSTTAVVTNPK.[E] |
| | | [R].YTIAALLSPYSYSTTAVVTNPKE.[-] |

**Supplementary Table 3.** Proportion of ATTRwt content in fibrils extracted from cardiac tissue.

| Sample | Percent ATTRwt (%) | Percent ATTRv (%) |
|---|---|---|
| ATTRv-P24S | 48.7 | 51.3 |
| ATTRv-A25S | 62.3 | 37.7 |
| ATTRv-D38A | 64.9 | 35.1 |

**Supplementary Table 4.** Root mean square deviation (RMSD of Cα atoms) of ATTRv and ATTRwt structures against their average consensus RMSD calculated by GESAMT.

| Structures | | RMSD (Å) |
|---|---|---|
| ATTRv-P24S | PDB: 8e7i | 0.586 |
| ATTRv-D38A | PDB: 9nnn | 0.639 |
| ATTRwt | PDB: 8gbr | 0.471 |
| ATTRwt | PDB: 8e7d | 0.430 |
| ATTRwt | PDB: 8e7h | 0.436 |
| ATTRwt | PDB: 8ade | 0.539 |

**Supplementary Table 5.** Complete data collection and refinement statistics.

| | **ATTRv-P24S** | **ATTRv-D38A** | **ATTRv-A25S** |
|---|---|---|---|
| PDB ID | 8e7i | 9nnn | n/a |
| EMD ID | 26692 | 49578 | 49581 |
| **Data collection** | | | |
| Microscope | Titan Krios | Titan Krios | Titan Krios |
| Voltage (kV) | 300 | 300 | 300 |
| Detector | K3 | K3 | K3 |
| Magnification | 81,000x | 105,000 | 105,000 |
| Software | EPU 3.5 | SerialEM | SerialEM |
| Total dose (e-) | 50 | 50 | 51 |
| Exposure time (sec) | 3 | 4 | 4.5 |
| Defocus range (μm) | -1.5 to -2.1 | -1.1 to -2.2 | -0.9 to -2.2 |
| No. of frames | 33 | 50 | 50 |
| Pixel size (Å) | 1.1 | 0.83 | 0.83 |
| **Reconstruction** | | | |
| Micrograph selected | 4,403 | 4,467 | 1288 |
| Software | Relion 3.1 | Relion 4.0 | Relion 3.1 |
| Box size (pixel) | 256 | 256 | 256 |
| Total extracted segments | 430,928 | 442,908 | 333,009 |
| Number of segments after 2D | 259,091 | 114,578 | 164,492 |
| Number of segments after 3D | 167,649 | 36,014 | 64,516 |
| Symmetry imposed | C1 | C1 | C1 |
| Helical rise (Å) | 4.93 | 4.74 | 4.75 |
| Helical twist (°) | -1.37 | -1.25 | -1.25 |
| Crossover length (Å) | 658 | 682 | 684 |
| B factor | -111.3 | -92.2 | n/a |
| Map resolution (Å; FSC=0.143) | 3.27 | 3.57 | 4.2 |
| Map resolution (Å; FSC=0.5) | 3.63 | 4.0 | 4.4 |
| **Model building and refinement** | | | |
| Non-hydrogen atoms | 3,585 | 3,570 | n/a |
| Protein residues | 460 | 455 | n/a |
| Number of chains | 5 | 5 | n/a |
| Water/ligands | 0 | 0 | n/a |
| MolProbity score | 1.87 | 2.12 | n/a |
| Clash score | 7.96 | 18.79 | n/a |
| Rotamer outliers (%) | 0 | 0 | n/a |
| R.M.S deviations bonds (Å) | 0.005 | 0.005 | n/a |
| R.M.S deviations angle (°) | 0.635 | 0.667 | n/a |
| Ramachandran plot<br>Favored<br>Allowed<br>Outliers | <br>93.18<br>6.92<br>0.00 | <br>95.4<br>4.60<br>0.00 | n/a |
| CaBLAM outliers (%) | 4.76 | 3.61 | n/a |
| Model vs Data | 0.84 | 0.85 | n/a |

**Supplementary References**


1. T. Uemichi, M. A. Gertz and M. D. Benson, *J Med Genet*, 1995, **32**, 279-281.
2. M. Yazaki, T. Yamashita, J. C. Kincaid, J. R. Scott, R. G. Auger, P. J. Dyck and M. D. Benson, *Muscle Nerve*, 2002, **25**, 244-250.
3. H. J. Cho, J. Y. Yoon, M. H. Bae, J. H. Lee, D. H. Yang, H. S. Park, Y. Cho, S. C. Chae and J. E. Jun, *J Cardiovasc Ultrasound*, 2012, **20**, 209-212.
4. M. Kishikawa, T. Nakanishi, A. Miyazaki, A. Shimizu, H. Kusaka, M. Fukui and T. Nishiue, *Amyloid*, 1999, **6**, 278-281.
